# Cytosolic Ca^2+^ as a universal signal for rapid root growth regulation

**DOI:** 10.1101/2025.10.17.683082

**Authors:** Marek Randuch, Ivan Kulich, Dmitrii Vladimirtsev, Shouguang Huang, Rainer Hedrich, Jiří Friml

## Abstract

Roots must continuously adapt their growth and have evolved the ability to rapidly respond to diverse environmental and hormonal cues, including the phytohormone auxin. Auxin-induced cellular responses involve membrane depolarization, cytosolic calcium (Ca²⁺) elevation, and extracellular pH increase, but how these processes lead to rapid growth regulation remains unclear. Here, we show that cytosolic Ca²⁺ acts as a universal signal integrating diverse cues for rapid regulation of root growth. Using live imaging, microfluidics, and optogenetics in Arabidopsis, we demonstrate that a swift rise in cytosolic Ca²⁺ is the primary target of auxin signaling affecting growth inhibition. Disruption of Ca²⁺ influx abolishes these responses, whereas light-gated Ca²⁺ influx from the apoplast or endoplasmic reticulum stores inhibits growth. Multiple unrelated stimuli—including auxin, extracellular ATP, RALF peptides, and hydrogen peroxide—converge on this Ca²⁺-dependent mechanism. Cytosolic Ca²⁺ elevation thus represents a necessary and sufficient step for rapid growth inhibition, revealing a unifying principle of root signaling.

## INTRODUCTION

Roots of land plants face the challenging task of navigating a highly complex soil environment filled with various obstacles and patches differing in humidity and nutrient content, as well as with herbivores, pathogens, and symbionts. In addition, roots perceive the status of aerial tissues via endogenous signaling. The ability to efficiently perceive, integrate, and respond to all these signals on very short timescales represents a strong evolutionary advantage (*1*, *2*). Indeed, much of the research in the past decades has shown that root growth is influenced by dozens, or perhaps hundreds, of different signals; some of them, such as the plant hormone auxin, can elicit cellular responses within seconds (*3*).

Whereas sustained developmental responses have been the target of various genetic screens in past decades - and the corresponding signaling mechanisms for transcriptional reprogramming have been elucidated (*2*) - the rapid responses remain much more enigmatic. Only recent advances in live imaging (*4*), root microfluidics (*5*), and optogenetic manipulation of cellular processes in plants (*6–10*) have made these rapid responses accessible to experimental investigation.

Auxin is a major regulator of root development, with a well-documented role in determining root architecture (*2*, *11*) and gravitropism (*12*). It mediates both transcriptional reprogramming responses that depend on nuclear auxin signaling (*11*), and very rapid responses (*13*) that rely, at least in part, on cell surface (*14*, *15*) and cytosolic (*16–18*) signal transduction mechanisms. Root growth inhibition is accompanied by a rapid cytosolic Ca²⁺ increase (*13*), plasma membrane (PM) depolarization (*18*), apoplast alkalinization (*14*) and apoplastic ROS production. While apoplastic ROS does not appear to be directly involved in growth regulation (*19*), apoplast alkalinization is highly relevant, as it has been proposed as an important integrator of signaling (*20*). According to the classical acid growth theory, apoplast alkalinization is equivalent to the drop of proton-motive-force, which is driving the H^+^-coupled solute transport, resulting in a decrease in turgor. Additionally, cell wall loosening enzymes are less active in a more alkaline pH leading to stiffening of the cell wall; both these processes inhibiting growth (*21*). Also, auxin-triggered cytosolic Ca²⁺ signaling is required for rapid growth inhibition, since mutations in the CYCLIC NUCLEOTIDE-GATED CHANNEL14 (CNGC14) Ca²⁺ channel (*13*, *22*) and in ARMADILLO REPEAT ONLY (ARO) CNGC regulators (*23*), both show defects in this process.

Besides auxin, some other hormonal and non-hormonal signals have been associated with rapid growth regulation and many more with the cytosolic Ca²⁺ increase. For example, the RAPID ALKALINIZATION FACTOR (RALF) secreted peptide involved in cell wall integrity monitoring, causes rapid growth arrest (*24*, *25*). On the other hand, extracellular glutamate (*26*) and ATP (eATP), that act as damage-associated molecular patterns, trigger increases in cytosolic Ca²⁺ (*27*). Other examples include hydrogen peroxide (H₂O₂) (*28*), as well as mechanical and cold stress (*29*, *30*); all have been shown to elevate cytosolic Ca²⁺. Thus, many signals can regulate cytosolic Ca²⁺ levels, either through various CNGC or other type of Ca²⁺ channels. Nonetheless, whether these Ca²⁺ activations are linked to growth regulation - analogous to auxin - and what the underlying mechanisms are, remain unclear.

Here, we show that an increase in cytosolic Ca²⁺ is the primary target of auxin signaling and represents the necessary and sufficient step for rapid root growth inhibition. Consequently, multiple signals, both endogenous and exogenous, converge on the regulation of cytosolic Ca²⁺ as a universal mechanism for rapid growth regulation.

## RESULTS

### Timing of cellular auxin responses during rapid root growth regulation

As shown previously (*11*), rapid, auxin-triggered root growth inhibition is accompanied by rapid cellular responses including cytosolic Ca^2+^ increase, membrane depolarization and extracellular pH alkalinisation (Fig. 1A-E; fig. S1). To gain insight into the hierarchy of these events, we analysed their timing following auxin (IAA) application.

**Fig. 1.**
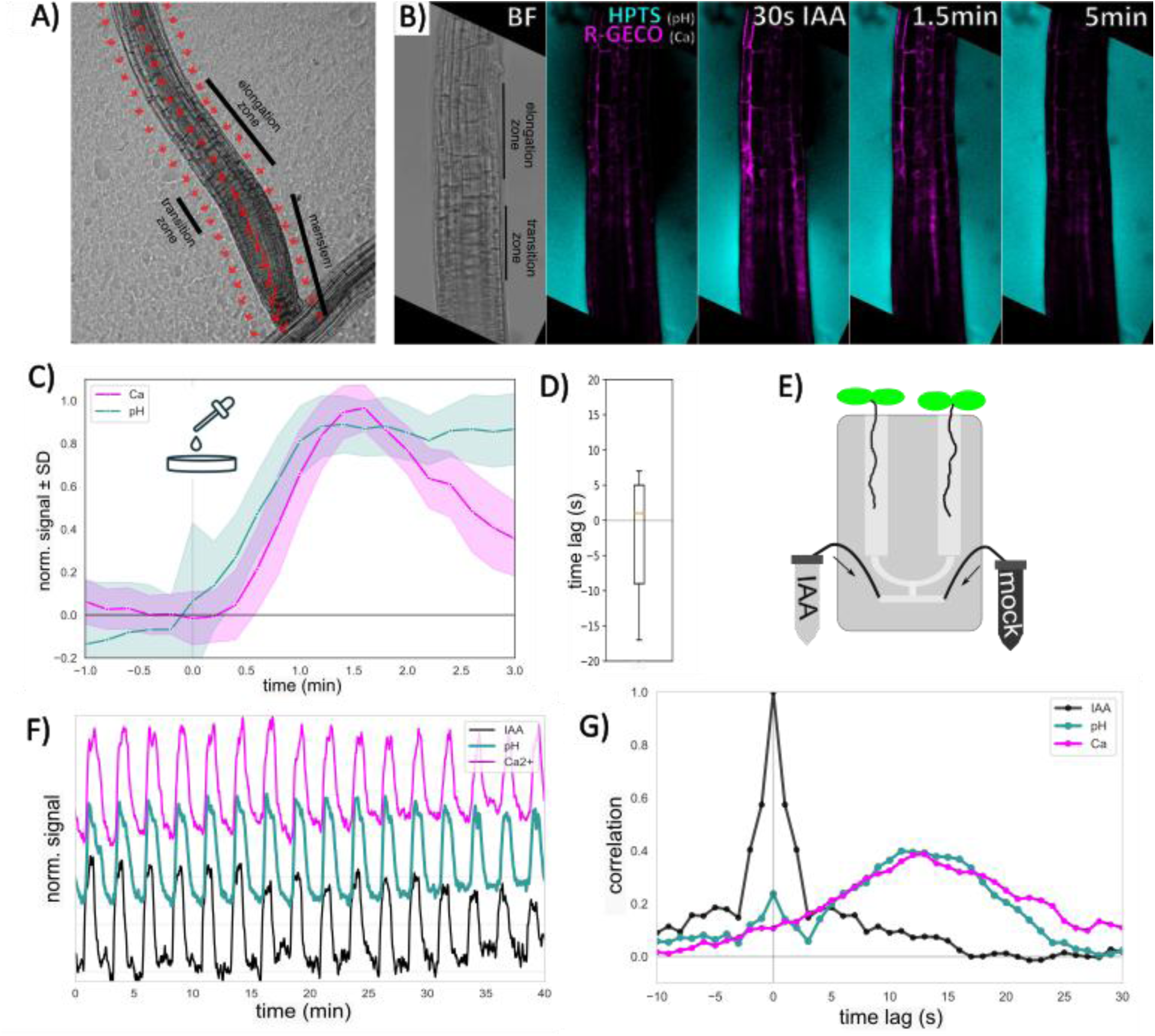
Rapid dynamics of auxin-induced rise in cytosolic Ca^2+^ and apoplastic pH. **(A)** The zonation of the *Arabidopsis* root, with arrows illustrating a growth shift between two timeframes. **(B)** The acidity of elongation zone is manifested by low intensity HPTS_470nm_ halo (pH, cyan), the transition zone has intense (basic) halo before treatment. After 30 s of IAA treatment: visible global rise of R-GECO signal (Ca^2+^, magenta) and burst of HPTS signal (pH) in transition zone; After 1.5 min: HPTS signal plateaus in elongation zone. **(C)** HPTS halo and R-GECO respond similarly in elongation zone to sharp (<1 s, fig. S1E) application of 100 nM IAA. **(D)** The temporal lag determined by cross-correlation analysis of signal pairs shown in (D; n=7, T-test p=0.51) **(E)** Design of 2-inlet RootChip allowing medium exchange under constant total flow. **(F)** IAA oscillations in RootChip are inducing R-GECO and HPTS responses, consistently over time. **(G)** The cross-correlation of R-GECO and HPTS signal derivatives with positive derivative of AF^647^ tracking dye signal (AF647′ if AF647′ > 0, else 0)), isolating IAA injections while excluding washout phases. The time of the curve maximum represents lag of response after IAA treatment.

Typically, we applied 100 nM IAA (fig. S1E) to 4-days-old seedlings in the medium under the microscope. We visualized cytosolic Ca^2+^ using GCaMP and R-GECO fluorescent sensors (*26*, *31*), which respond with a similar dynamic (fig. S2). For the extracellular pH we used ratiometric fluorescent dye HPTS (*32*) and for the membrane potential, the DiSBAC_2_(3) dye and impalement electrode (*16*, *18*).

The GCaMP revealed that Ca^2+^ spike initiates in the root transition zone after about 10 s, the increases then spreads in both directions to elongation and meristematic zones, peaking in 2 min and starting to decrease in the whole root tip after 5 min (Fig. 1B,C; fig. S1). HPTS showed a similar, initial burst of apoplastic alkalinisation in the transition zone (fig. S1) with a decrease after 5 min similarly to GCaMP, and a slower build-up in the elongation zone where the alkalinisation persists even after partial decrease of Ca^2+^ (Fig. 1B,C). Simultaneous imaging of R-GECO with HPTS confirmed a similar dynamic of initiation in the elongation zone (Fig. 1C,D; fig. S3; Movies S1). On the other hand, the membrane potential measurements using DiSBAC showed a much-delayed response (fig. S4,5; sup. Movies S2); therefore, we focused in our further analysis predominantly on timing of Ca^2+^ and pH responses.

Our single-treatment observations would suggest a slightly faster onset of pH response before the cytosolic Ca^2+^ increase in the elongation zone (Fig. 1C), nonetheless, given the slightly different response dynamics of used tools, this approach could not conclusively distinguish timing of these responses. Therefore, we used oscillations of IAA treatments in RootChip microfluidics (*19*) (Fig. 1E,F) and cross-correlations of the response derivations (Fig. 1G). Even this approach was not able to temporally distinguish these responses; confirming that both, cytosolic Ca^2+^ and apoplastic pH responses occur with highly similar, rapid dynamics of about 10 s.

Overall, these observations consistently show that the rapid cellular responses to auxin are first visible in the transition zone with cytosolic Ca^2+^ spike and extracellular pH alkalinisation showing fast initiation.

### Apoplast alkalinization but not PM depolarization is sufficient for growth inhibition

The timing of events suggests that PM depolarization occurs as a later step in growth regulation. (fig. S4). To further test whether depolarization is a more downstream response, we utilized a RootChip to manipulate the PM potential by extracellular K^+^/NH_4_^+^ exchanges. By replacing the AM medium containing 20 mM K⁺ by 20 mM NH_4_^+^, we repeatedly hyperpolarized and depolarized the PM by approximately 40 mV as confirmed by intracellular voltage-recording electrodes (Fig. 2A). K^+^-induced depolarisation lead only to minor (<15%) and transient root growth inhibition as compared to much stronger (>50%) and sustained inhibition by IAA (fig. S1F). Immediately after K⁺-induced depolarisation, we observed no increase (Fig. 2B). These observations suggest that PM depolarisation itself is insufficient to mediate cytosolic Ca^2+^ increase and growth inhibition.

**Fig. 2.**
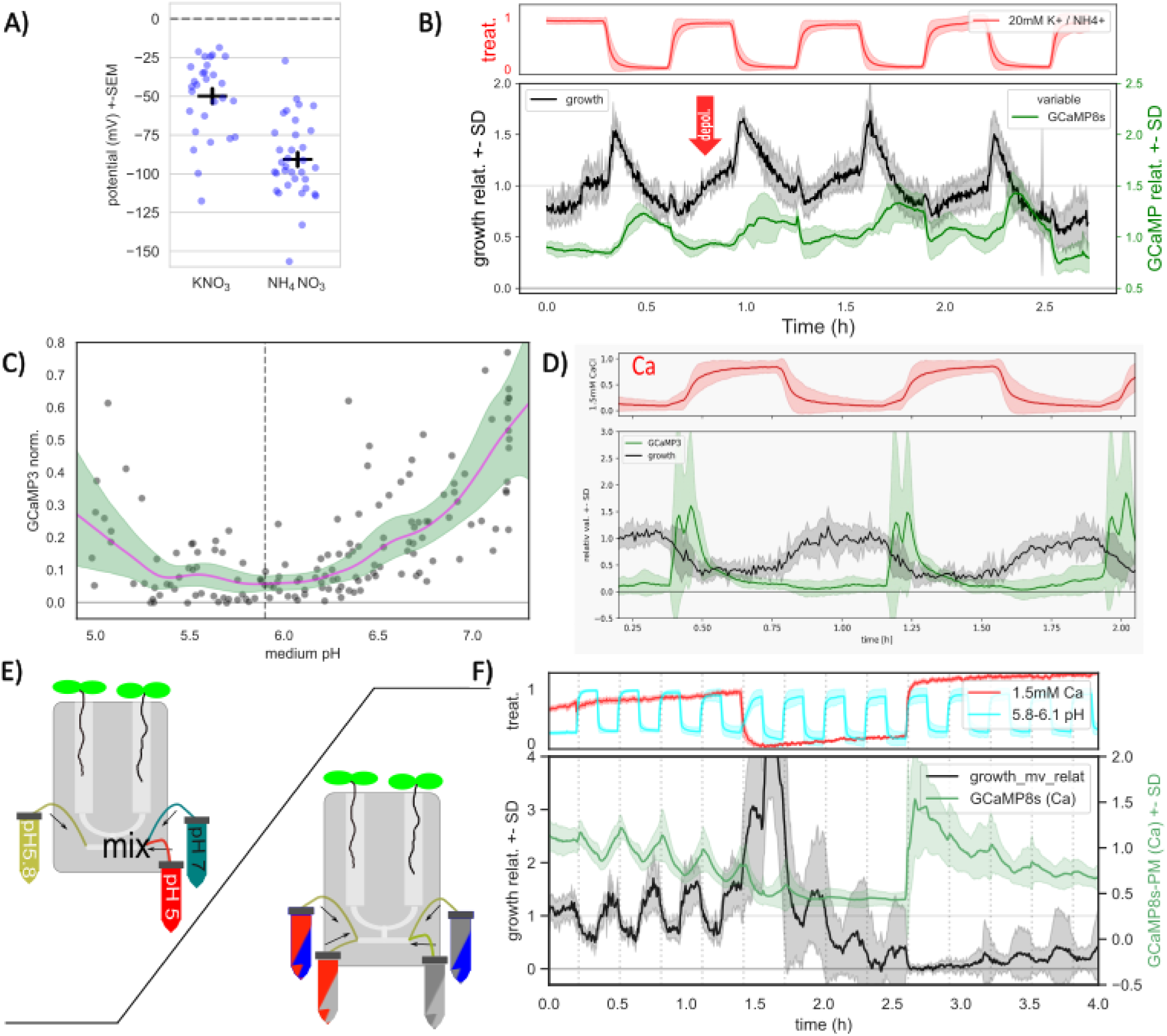
PM polarization, apoplastic pH and cytosolic Ca^2+^ in root growth regulation. **(A)** Replacement of 20 mM NH_4_^+^ by 20 mM K^+^ in medium cause depolarisation as measured by impalement electrode. (n>29, Wald test p<0.001, 2 repetitions combined) **(B)** 20 mM NH_4_^+^ to K^+^ exchange (red arrow) is not causing root growth (black) imbibition or increase in GCaMP signal (Ca^2+^, green). (representative from 3 repetitions) **(C)** Each point represent cycle of medium pH change from 5.8 to more acidic or alkaline pH. Average rise of GCaMP signal is plotted. (details in sup. fig 6.) (2 experiments merged) **(D)** Washing roots with Ca^2+^-free medium is causing drop of GCaMP signal and acceleration of growth while supplementing Ca^2+^ is causing growth inhibition. **(E)** Design of RootChip for exchanging mock and mixture of two stock solutions (used in C), 4-inlet RootChip used in (D). **(F)** Medium alkalinisation (blue down) is inhibiting growth accompanied by rise of GCaMP on Ca^2+^-containing medium (red up). On Ca^2+^-free, 1 mM BAPTA supplemented medium (red down) alkaline medium is still inhibiting growth without GCaMP response. (representative from 2 repetitions)

As we were unable to distinguish the rapid timing of cytosolic Ca^2+^ spike and extracellular pH alkalinisation, to gain insight into hierarchy of these events, we tested their interdependence in terms of root growth regulation. By manipulating the RootChip medium pH from 5.8 up to 1 pH units in both directions (Fig. 2C,E), we observed that both alkalinisation and acidification caused an increase in the GCaMP-monitored cytosolic Ca^2+^ (Fig. 2C, fig. S6) with a steeper GCaMP increase for alkalinisation (pH > 6.3). This suggests that either Ca^2+^ acts downstream of pH changes, or rather, that there is a feed-back regulation between these processes consistent with previous works showing that manipulating H^+^ fluxes impacts cytosolic Ca^2+^ (*7*).

Considering previous observations that apoplastic pH manipulations influence root growth rates (*14*), we tested whether extracellular pH alkalinization alone can inhibit root growth also in the absence of cytosolic Ca^2+^ increase. Using a microfluidic 4-inlet setup enabling application of two mock/treatments in all four combinations (Fig. 2E), we confirmed that exchanging the medium with pH 5.8 to pH 6.1 caused growth inhibition, which was accompanied by initial increase in cytosolic Ca^2+^ followed by a rapid drop (Fig. 2F). The same pH manipulations performed in the nominally Ca^2+^-free media, supplemented by 1 mM BAPTA, did not results in any cytosolic Ca^2+^ increase but the root growth was still inhibited as in the standard medium (Fig. 2F). This suggests that apoplastic pH is sufficient to regulate the root growth even in absence of the cytosolic Ca^2+^ increase.

These observations show that PM depolarization is not sufficient to inhibit growth, whereas apoplast alkalinization is. Notably, this occurs also in the absence of cytosolic Ca^2+^ spike, implying that apoplastic pH likely acts as effector downstream of cytosolic Ca^2+^.

### Many signals converge on cytosolic Ca^2+^ to regulate root growth

The notion that cytosolic Ca^2+^ acts upstream of pH regulation for root growth regulation aligns with the published findings that *cngc14* mutations affecting CNGC14 Ca^2+^ channel leads to auxin-insensitive root growth (*13*, *22*). We confirmed that the rapid growth inhibition, apoplast alkalization, and PM depolarization induced by 100 nM IAA are abolished in the *cngc14* roots (fig. S7). Similarly, in the *aro2/3/4* mutant, which is defective in function of presumably all CNGC channels (*23*), these auxin responses are abolished (Fig. 3C).

**Fig. 3.**
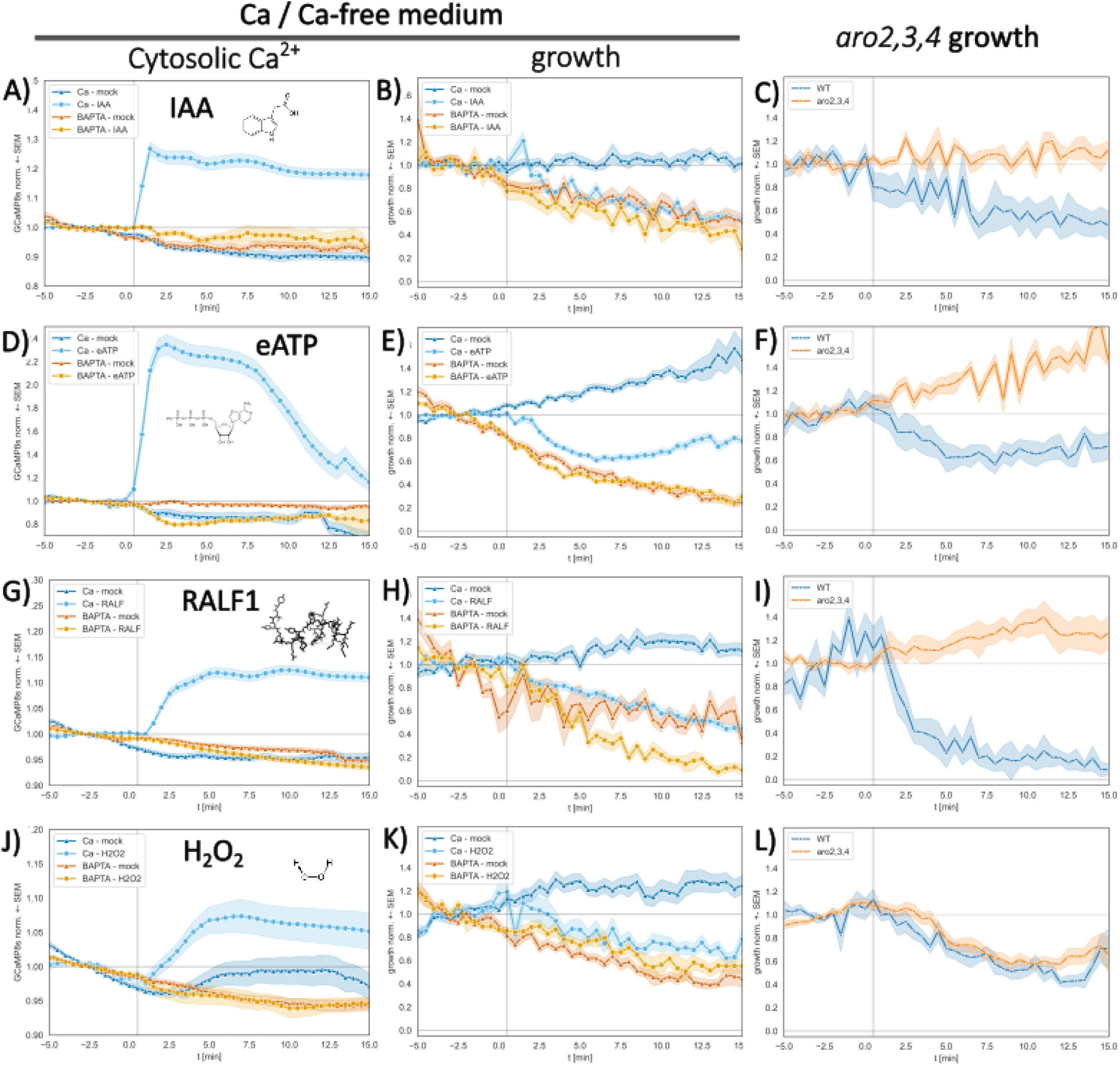
Cytosolic Ca^2+^ increase as common signal for growth inhibition by diverse stimuli. Treatment with 100 nM IAA **(A-C)**, 100 µM eATP **(D-F)**, 1 µM RALF peptide **(G-I)** and 200 µM H_2_O_2_ **(J-L)** is causing the GCaMP signal increase on Ca^2+^-containing medium (A,D,G,J), but not on 0.1 mM BAPTA Ca^2+^-free medium (B,E,H,K). All treatments are rapidly inhibiting growth on Ca^2+^-containing medium. However, 0.1 mM BAPTA-containing, Ca^2+^-free medium prevent or delay this growth inhibition (C,F,I) (n>20, combined 3 replicates). The growth of *aro2/3/4* mutant is insensitive to IAA, eATP and RALF (L) (n>6, combined 2 replicates), while H_2_O_2_ induces the same growth response as mock (n>18, combined 3 replicates).

To verify these genetic approaches, we established monitoring rapid root growth responses using Ca^2+^-free medium (*19*) supplemented with 0.1 mM BAPTA. This leads to an abolishment of rapid auxin responses including cytosolic Ca^2+^ increase and root growth inhibition (Fig. 3A,B). These observations confirm that auxin-induced cytosolic Ca^2+^ increase acts upstream of root growth inhibition.

Next, we tested whether this Ca^2+^-dependent growth regulation is specific to auxin or can occur also downstream of other known Ca^2+^ signal-provoking physiological effectors. To answer this question, we tested three signals known to induce Ca^2+^ increase in roots: extracellular ATP (eATP) – small molecule functioning as cell damage signal (*27*), RAPID ALKALINISATION FACTOR 1 (RALF) - peptide involved in monitoring cell wall integrity (*24*, *25*) and H_2_O_2_ - multirole signaling/defense compound (*19*, *28*, *33*). All three treatments increase GCaMP signal in roots (Fig. 3D,G,J), confirming that they indeed induce cytosolic Ca^2+^ increase.

When we treated roots with eATP, RALF and H_2_O_2_ in the Ca^2+^-free medium, the GCaMP signal increase did not occur (Fig. 3D,G,J). This confirms that for all these signals, as shown for auxin, Ca^2+^ influx from the apoplast is the sole contributor to the cytosolic Ca^2+^ increase. This allowed us to test the importance of cytosolic Ca^2+^ for the growth regulation downstream of these signals. All three effectors inhibited rapidly root growth in standard medium, whereas this response was delayed or completely abolished in the Ca^2+^-free medium (Fig. 3E,H,K).

These findings we confirmed genetically, taking advantage of the *aro2/3/4* mutant affected generally in the CNGC function (*23*). eATP- and RALF-induced root growth inhibition was absent in *aro2/3/4*, whereas H_2_O_2_ was still effective (Fig. 3F, I, J), suggesting that H_2_O_2_-induced Ca^2+^increase does not require CNGC function but other types of Ca^2+^ channels or transporters (*34*).

These pharmacological and genetic studies revealed that Ca^2+^ influx from the apoplast and resulting cytosolic Ca^2+^ increase is essential not only for the auxin-induced root growth inhibition, but also acts downstream of signals rapidly inhibiting root growth, as exemplified by IAA, eATP, RALF and H_2_O_2_.

### Activation of light-gated Ca^2+^ channels at the PM or ER is sufficient to regulate growth

To test whether cytosolic Ca^2+^ is not only required but also sufficient to inhibit growth, we used microfluidics and replaced 1.5 mM CaCl_2_ by 1.5 mM MgCl_2_. Reducing the Ca^2+^ levels in the medium caused accelerated root growth while re-supplementing Ca^2+^ inhibited the growth (Fig. 2D). Further Ca^2+^ depletion using 1 mM BAPTA led to a 6-fold increase in growth rate (Fig. 2F) and rupture of root hairs (Movies S3), implying that Ca^2+^ functions as a constant growth “break” also under standard conditions. Restoring the 1.5 mM CaCl_2_ in the medium resulted in cytoplasmic Ca^2+^ transients and ultimately to growth inhibition (Fig. 2D,F). This suggests that manipulation of Ca^2+^ levels is sufficient to induce rapid responses leading to the root growth inhibition.

To test specifically whether the cytosolic Ca^2+^ transients are the sufficient signal to induce responses leading to root growth inhibition, we established optogenetic experiments in Arabidopsis roots and utilized the blue light-activated cation channel XXM2.0, which has been described as a tool for Ca²⁺ manipulation in plants (*6*, *35*) (Fig. 4A,B). During 5 min blue light illumination (200 µmol·s^-1^·m^-2^), XXM2.0 roots exhibited pronounced increased in R-GECO1 signal, indicative of a rise in cytosolic Ca^2+^, whereas the response in wild-type (WT) roots was minimal (Fig. 4C). The apoplastic pH reported by HPTS in light-treated (488 nm excitation light for HPTS imaging served also for XXM activation) XXM2.0 roots followed a pattern similar to that observed in IAA-treated roots, including an alkalinisation burst in the transition zone (Fig. 4D,E). Blue light treatment also triggered reversible rapid root growth inhibition in XXM2.0 roots, with growth rates gradually returning to pre-stimulus level, once the light was turned off (Fig. 4F). These findings confirm that opening of XXM2.0 channels and resulting influx of Ca^2+^ ions to the cytosol is sufficient to replicate the rapid responses typically seen following treatment with IAA.

**Fig. 4.**
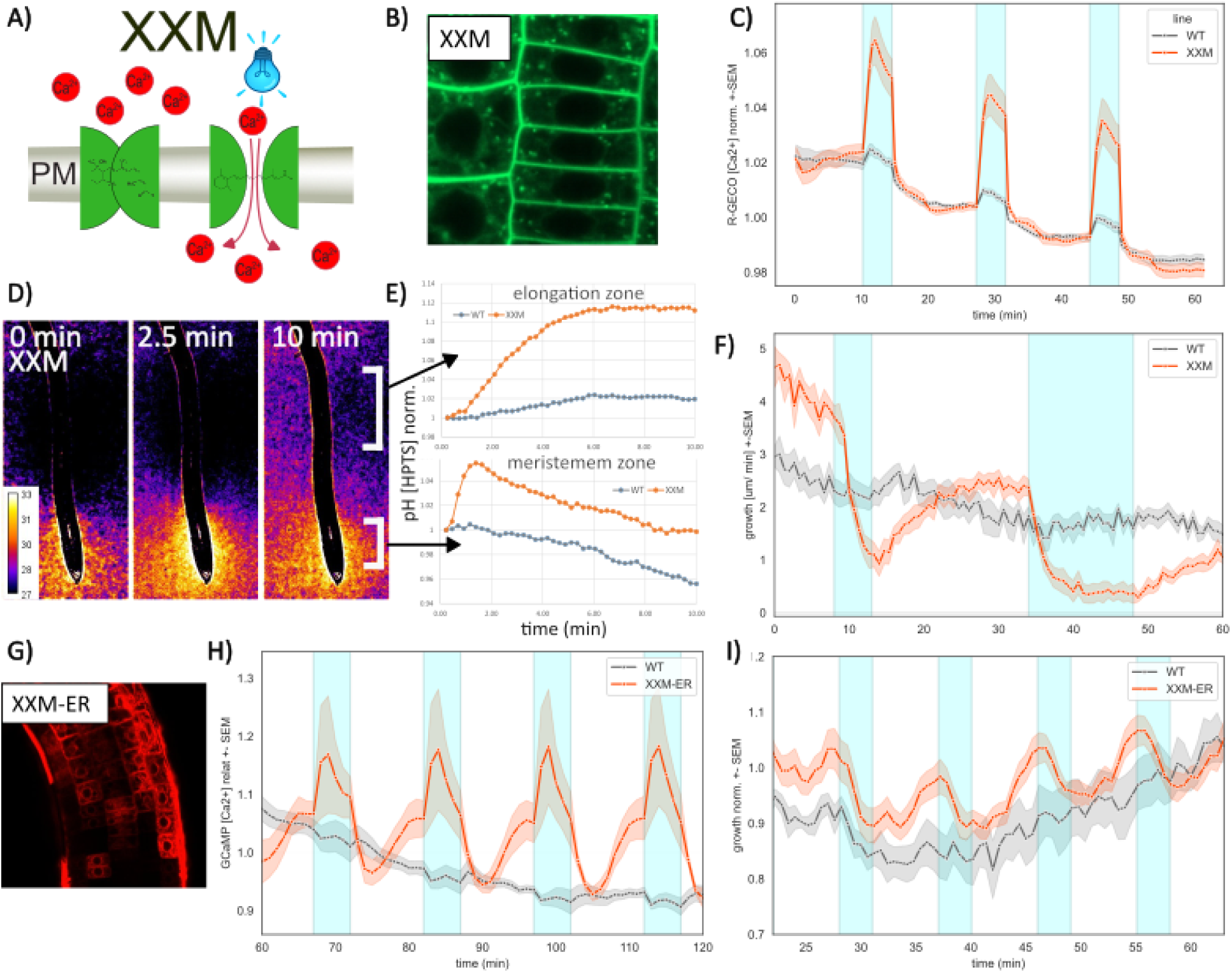
Optogenetically induced cytosolic Ca^2+^ in apoplast pH and growth regulation. **(A)** Schematics of light-gated cation channel XXM-YFP. **(B)** XXM-YFP localization to the PM. **(C)** Blue light-induced cytosolic Ca^2+^ increase as reported by R-GECO. (representative from 2 replicates) **(D-F)** Blue light-induced HPTS-reported apoplast alkalinisation (D,E) and rapid and reversible growth inhibition (F) in XXM2.0 but not WT roots (n>7, representative from 3 replicates). **(G-I)** XXM-ER line expressing ER-targeted XXM-mCherry (G) but not WT responds to blue light by the Ca^2+^ increase as reported by R-GECO (H) and by rapid and reversible growth inhibition (I) (n>8, 2 replicates).

Nonetheless, XXM, despite being primarily used as a Ca^2+^ channel, may allow also for the passage of H^+^ (*36*). Thus, it cannot be fully excluded that the H^+^ leakage from the apoplast might be partly responsible for the observed effects. To address these issues, we created an XXM-mCherry version and targeted it to the ER membrane (XXM-ER) (Fig. 4G). The ER contains high Ca^2+^ levels; however, under normal conditions, the pH inside the ER lumen is similar to that of the cytoplasm (*37*). After blue light exposure, XXM-ER but not WT roots showed an increase in the cytosolic Ca^2+^ as reported by GCaMP (Fig. 4H). Also, the XXM-ER line displayed rapid root growth inhibition in response to blue light, while WT roots exhibited constant growth (Fig. 4I). These results further solidify the notion that an increase in cytosolic Ca^2+^, regardless of the Ca^2+^ source involved, is the key for growth inhibition.

This shows that the cytosolic Ca^2+^ increase is not only necessary but also sufficient to induce rapid root growth inhibition.

## DISCUSSION

In recent years, a set of different signals has been identified for causing rapid root growth regulation, among them the plant hormone auxin is of prime importance. Auxin-induced root growth inhibition is accompanied by apoplast alkalinization, an increase in cytosolic Ca²⁺, and plasma membrane depolarization. Such rapid, non-transcriptional effects, as opposed to sustained developmental responses, have been difficult to track genetically, and thus the underlying mechanisms remain comparatively obscure. Recent advances in live imaging (*4*), root microfluidics (*5*, *17*) and optogenetics (*6–10*, *35*), which allow immediate control and monitoring of cellular processes, open new possibilities for gaining mechanistic insights into these regulations.

### Cytosolic Ca²⁺ increase is the primary auxin response upstream of growth regulation

To gain insight into the causal hierarchy of the rapid cellular events downstream of auxin, we examined their timing and manipulated each of these processes separately. Apoplast alkalinization and cytosolic Ca²⁺ increase occur with very similar dynamics, initiating about 10 seconds after the auxin signal. In contrast, membrane depolarization occurs relatively late and is not sufficient to regulate growth (Fig. 2B; fig. S4), suggesting that depolarization is rather downstream of the early auxin response.

Manipulation of apoplastic pH and cytosolic Ca²⁺ revealed that they represent very early, interdependent events, both required for root growth regulation. An increase in cytosolic Ca²⁺ leads to apoplast alkalinization (Fig. 4D) and, conversely, apoplast alkalinization induces a cytosolic Ca²⁺ transient (Fig. 2C), implying tight feedback regulation. Nonetheless, apoplast alkalinization does not occur when the cytosolic Ca²⁺ increase is prevented, either genetically or pharmacologically (Fig. 3). On the other hand, apoplastic pH impacts growth even when the cytosolic Ca²⁺ transients are prevented (Fig. 2F). This places cytosolic Ca²⁺ as the primary cellular auxin response upstream of apoplastic pH regulation and root growth regulation.

### Cytosolic Ca^2+^ is a necessary and sufficient signal for root growth regulation

Preventing Ca²⁺ transients, either by placing roots in Ca²⁺-free medium or by genetically interfering with the function of CNGC Ca²⁺ channels, prevents apoplast alkalinization and root growth inhibition (Fig. 3; fig. S7) (*13*, *22*). This strongly suggests that cytosolic Ca²⁺ transients are a necessary component of root growth regulation by auxin.

Observing the growth of roots and root hairs under Ca²⁺-deficient or Ca²⁺-supplemented conditions (Fig. 2D,F) suggests that Ca²⁺ acts as a “brake,” dynamically regulating growth under physiological conditions—preventing overextension and bursting of root hairs (*38*) and poising the primary root for both positive and negative dynamic regulation.

Manipulating Ca²⁺ levels in the medium or optogenetically opening Ca²⁺ channels to allow passage of Ca²⁺ from the apoplast to the cytosol are both sufficient to induce downstream apoplast alkalinization and root growth inhibition (Fig. 4D-F). This not only confirms that Ca²⁺ transients act upstream of apoplastic pH regulation but also shows that they are a sufficient signal for root growth regulation. This is further supported by experiments using an optogenetically controlled Ca²⁺ channel at the ER membrane, where light-induced release of Ca²⁺ from ER stores into the cytosol is likewise sufficient to inhibit root growth (Fig. 4I).

### Multiple signals converge on cytosolic Ca^2+^ to regulate root growth

Given, that cytosolic Ca²⁺ increase, regardless of its source, is the sole, sufficient signal for root growth inhibition, it may also serve as a central mechanism through which other signals, beyond auxin, control growth. To test this possibility, we selected several chemically and functionally diverse signals potentially linked to rapid growth regulation and/or cytosolic Ca²⁺ dynamics. These included extracellular eATP, RALF secreted peptide and H₂O. All these signals triggered a remarkably similar suite of cellular responses to auxin, including cytosolic Ca²⁺ transients, and subsequent inhibition of root growth.

By combining genetic and pharmacological interference with microfluidics and live-cell imaging, we found that, similar to auxin, these signals induce cytosolic Ca²⁺ transients that are required for root growth inhibition (Fig. 3). Notably, while auxin, eATP, and RALF promote Ca²⁺ influx through CNGC-type Ca²⁺ channels, H₂O₂ employs a distinct mechanism to elevate cytosolic Ca²⁺.

Together, these findings point to a shared mechanism of rapid growth regulation, in which cytosolic Ca²⁺ elevation represents a critical step downstream of multiple, diverse signalling pathways (Fig. 5). Such a universal regulatory module likely operates not only in hormonal signalling but also during responses to biotic and abiotic stress. Moreover, this work provides both a conceptual framework and an experimental toolbox for testing whether other environmental or endogenous cues utilize the same mechanism.

**Fig. 5.**
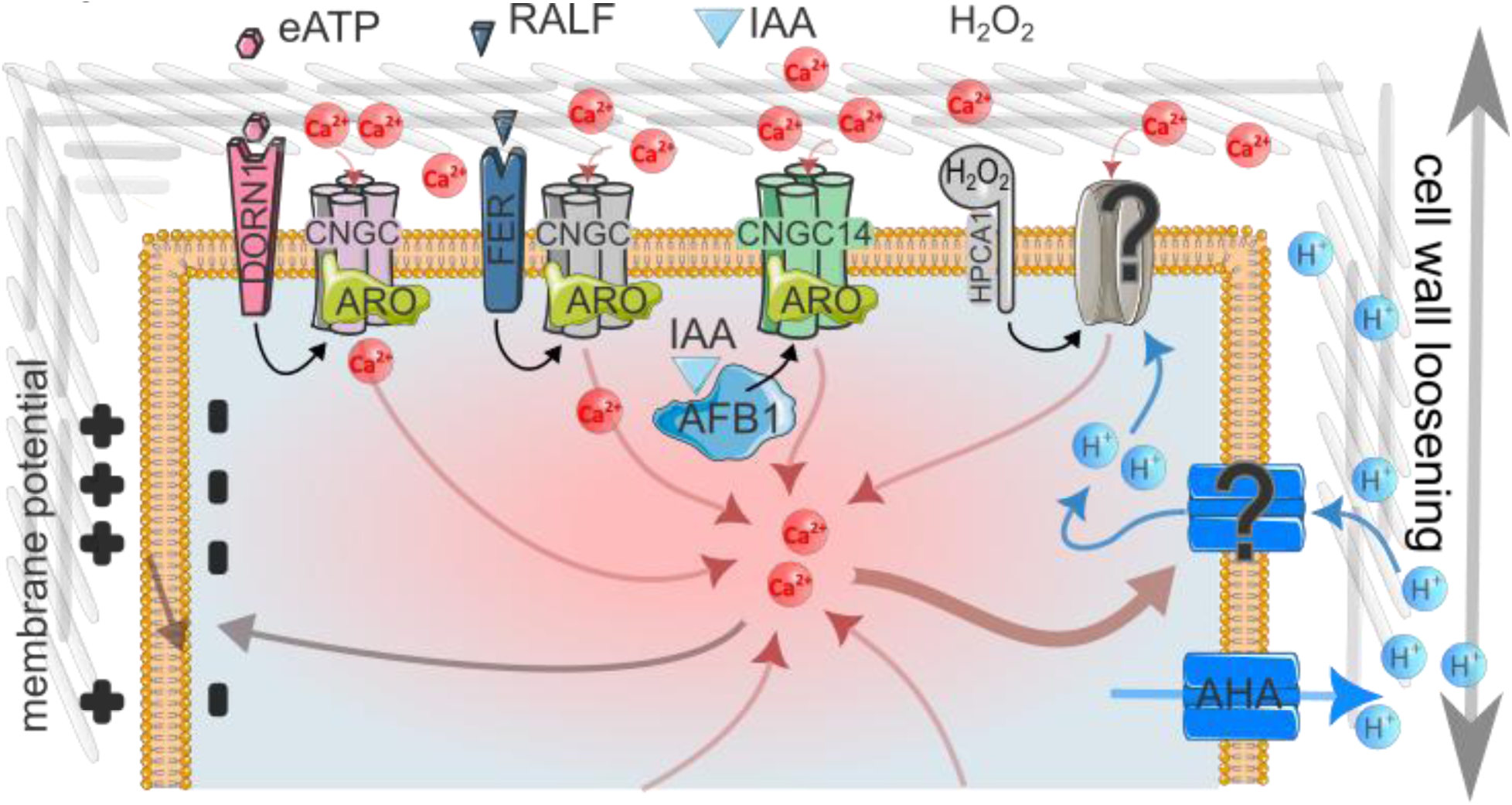
Diverse stimuli converge on cytosolic Ca^2+^ for growth regulatio. Diverse stimuli mediate rise in cytosolic Ca^2+^ via influx from the apoplast through CNGC-type channels (IAA, eATP, RALF) or other type of Ca^2+^ transport (H_2_O_2_). The cytosolic Ca^2+^ increase is sufficient for apoplast alkalinisation, leading to growth inhibition. Cytosolic Ca^2+^ and apoplastic pH is linked by a feedback loop. On the other hand, membrane depolarisation is acting downstream and is insufficient to inhibit growth alone. (Icons adapted from Servier Medical Art, CC BY 3.0).

## ACKNOWLEDGMENTS

We gratefully acknowledge Dr. Andrea Bleckmann, creator of pBLAX000 vector and Dr. Shiqiang Gao for donation of template used in XXM-ER cloning. This research was supported by the Laboratory Support Facility, the Plant Facility, the IT Facility and the Imaging and Optical Facility of the Institute of Science and Technology Austria. The graphical elements used in Fig. 5 were adapted and modified from Servier, https://smart.servier.com, and are used under CC BY 3.0 licence. The authors acknowledge the use of AI-tools (Microsoft Copilot, Gemini) for language refinement. All content was critically reviewed by authors.

## Author contributions

Conceptualization: JF, MR, IK

Methodology: MR, IK, DV

Investigation: MR, IK, DV, SH

Visualization: MR

Funding acquisition: JF, RH

Supervision: JF, RH

Writing – original draft: JF, MR

Writing – review & editing: JF, MR, IK, RH, DV, SH

## Data and materials availability

All data are available in the main text or the supplementary materials. For material requests contact JF.

## SUPPLEMENTARY MATERIALS

## Materials and Methods

### General growth condition

The all used Arabidopsis lines are in Columbia-0 (*Col-0*) background. The *afb1-3* line was obtained from (Savaldi-Goldstein et al., 2008), *cngc14-2* from (Shih et al., 2015), GCaMP3 from (Toyota et al., 2018), *R-GECO* from (Keinath et al., 2015) and *aro2/3/4* T-DNA insertion line was described previously (Kulich et al., 2020; Kulich, Oulehlová, et al., 2025) and *XXM2.0* was described in (Ding et al., 2024).

For growing and experiments was used standard Arabidopsis Medium (“AM+”): 1/2 MS salts (Duchefa, M0221), 1% sucrose (Sigma-Aldrich, S0389), 500mg/L MES (Duchefa, M1503), pH 5.8 adjusted by KOH (Merck, 105021), if not specified otherwise. The seeds were treated by - 20°C for 2 days and then surface sterilised by chlorine gas (∼16 h), plated on 1% agarose AM+ plates and stratified for 2 days at +4°C. Than grown on 16-h-light/8-h-dark light cycle and 21°C for 3-4 days.

### Direct application of treatments

To generate Fig. 1B-D and fig. S1, seedlings were prepared for live imaging by mounting in glass-bottom chamber. Seedlings were positioned on glass with stripe of double side tape, so the root tips are reaching out of the tape. The root tip (∼1 mm) was covered by block of solid agarose and rest of the root was mounted by room temperature 2% low melting agarose (Promega). After an approximately one-hour rest period, the solid agarose block was carefully removed by raiser, exposing the root tip. The chamber was then filled with liquid AM+ medium. A mock treatment was performed prior to the experiment to verify the stability of the mounting and allow the root to adapt to slight agitation. Treatment was done by rapid addition (<1 s) of 1/3 of the AM+ volume with 4x concentration of treatment. Imaging was conducted using a Leica Stellaris 5 microscope with an HC PL APO 20x water objective. The exact time of treatment on the record was determined by the by distortion of the medium surface, visible in bright-field channel.

The florescence channels (R-GECO, GCaMP, HPTS_405_, HPTS_470_, DiSBAC, TRITC) were quantified with the ImagJ (Fiji 2.15.0) software. The images were stabilised by registration with StackReg plugin. Average intensity of florescence in cortex for DiSBAC, epidermis and cortex for R-GECO and GCaMP and region corresponding to transition end elongation zone was quantified. The average of HPTS_405_ and HPTS_470_ signal was taken from medium regions adjacent to root (∼50 µm wide) and also more distant, outside the root halo, as a reference. The normalised HPTS ratio was calculated as (HPTS_470_ ^halo^ / HPTS_405_ ^halo^) / (HPTS_470_ ^distal^ / HPTS_405_ ^distal^).

### Growth rate measurement

Root growth was recorded using a Nikon Ti2E microscope equipped with Plan Fluor 10x objective for experiments depicted on Fig. 3 or LSM880 equipped with Plan-Apochromat 10x objective for experiments depicted on Fig. 2. The approximate root middle line was manually tracked from the start of differentiation zone to tip, and marked by ordered points on multiple time frames with ImageJ software. The subsequent analysis was performed by custom python script rune on HPC. The manually marked points were interpolated in XY and time. Regularly spaced square regions were defined along the interpolated root line. The overall shifts of the root in the regions (growth + bending + stage drift), was quantified by subpixel registration on the bright-field images. Root elongation was then calculated as the sum of the distance changes within pairs of adjacent regions, thus compensating for shifts by root bending and stage drifts. Moreover, the average intensity of fluorescence channels (GCaMP, TRITC) in the regions was quantified in the pipeline.

### Microfluidics

The core of microfluidic setup adapted from (N. B. C. Serre et al., 2021) was previously described in (Kulich, Vladimirtsev, et al., 2025). Seedlings were positioned to opened, 100 µm deep channel of PDMS chip and then covered with coverslip. The glass and PDMS chip were pressed together by screwing the chip holder forming “sandwich-like” microscope insert. Medium flow was regulated using an OB1 pressure controller (ElveFlow) coupled with a MSF2 flow sensor placed upstream of the chip. This formed a PI regulator feedback loop, operated via ESI software. For chip with channel cross-section 0.2 mm^2^, the flow rate was maintained at 5μL/min. For larger chips channel cross-section ∼1.5mm^2^, a flow rate of 20 μL/min was maintained using syringe pumps. Medium exchange was monitored by TRITC-dextran (Sigma-Aldrich, T1162) or AlexaFluor647-dextran (AF^647^; Invitrogen, D22914) tracking dye, depending on the fluorescent reporters being used.

For mixing of stock solution, the two tubing were connected by T-junction and between chip inlet and the T-junction was placed 5 cm of 1/32’’ ID tubing. The Root chip then can contain mixture of up to 3 solutions (during the transition phase) (see Fig. 2E, fig. S6A): AM+ (pH 5.8, no tracking dye) acidic AM+ (pH 5.0, labelled with TRITC) and alkaline AM+ (pH 7.3, labelled with AF647). The ratio of solutions in the mixture was determined from relative intensity of TRITC and AF^647^ tracking dyes. Then resulting pH of the corresponding MES buffers mixture was calculated using the Henderson–Hasselbalch relation.

For the cross-corelation analysis of IAA oscillation induced responses (Fig. 1), the obtained signals were corrected for long term drifts by scaling on its moving average (∼10min). To measure the Ca2+/pH lag only after IAA treatment, not influenced by IAA washout phase, the negative derivation of tracking dye signal (ΔAF^647^/Δt < 0), corresponding to IAA washout was set to be 0. The cross-corelation was done by Python scipy (1.11.4) package.

### Potential measurement with impalement electrode

Pulled borosilicate glass pipette (20-25 MΩ resistance), filled with 3 M KCl, was impaled by motorized system firs to epidermal cell approximately in transition zone and then to more proximal cells for repetitions. Recordings were made using a MultiClamp 700B amplifier and digitized by a Digidata 1550A at 10 Hz, the system was operated by Multiclamp and pCLAMP software. The amplifier was set to Current-Clamp mode. The records were randomised and blindly scored on potential maintained stably for at least 5 s. Measurements with evident technical issues were excluded. On the data pulled from two experiment, a linear mixed-effects model was fitted (statsmodels v0.13.5, REML) with treatment as a fixed effect, random intercepts for experiment and for plants nested within experiments. Significance of treatment effects was evaluated using Wald tests.

### Ca^2+^-free and K^+^-free medium

Stock solution was prepared from AM+ macronutrient components excluding CaCl_2_ (0.63mM KH_2_PO_4_ (Merck, 529568), 9.4 mM KNO_3_ (Sigma-Aldrich, P8291), 0.75 mM MgSO_4_ (Merck, 105886), 10.3 mM NH_4_NO_3_ (Sigma-Aldrich, A3795), 1% sucrose (Sigma-Aldrich, S0389) and 500 mg/L MES (Duchefa, M1503). Before use 0.1 mM BAPTA (Sigma-Aldrich 14510) was added to chelate Ca^2+^ µ-contamination and pH 5.8 was readjusted with KOH. Eider 1.5 mM MgCl_2_ (Sigma-Aldrich, 208337) or 1.5 mM CaCl_2_ (Sigma-Aldrich, C3306) was added for Ca^2+^-free or Ca^2+^-containing medium respectively. K^+^-free stock was prepared from 10.3 mM NH_4_NO_3_, 1.5 mM CaCl_2_ 0.63 mM NH_4_H_2_PO_4_ (Sigma-Aldrich, 09709), 0.75 mM MgSO_4_, 1% sucrose and 500 mg/L MES. pH 5.8 was adjusted with NaOH (Sigma-Aldrich, S5881). Eider 20 mM NH_4_NO_3_ or 20 mM KNO_3_ was added for K^+^-free or K^+^-containing medium respectively.

### Molecular cloning and plant transformation

XXM without Golgi transport signal was amplified by PCR (primers XXM-ER_toC & XXM-ER_toF) and cloned to BsaI (NEB, R3733L) linearised GreenGate vector pGGC000 (Lampropoulos et al., 2013) by NEBuilder (NEB, E2621X) according to manufacturer protocol. ER retention signal was inserted on C terminus of mCherry in pGGD003 block (Lampropoulos et al., 2013) by mutagenesis using PCR amplification (primers ERret_F & ERret_R) and NEBuider ligation. The following blocks were assembled to pGGZ001 (Lampropoulos et al., 2013) destination vector, modified to contain bacterial kanamycin resistance (pBLAX000): a) UBQ:XVE+LexA, b) N-decoy, c) XXM-ER, d) mCherry-ER, e) HSP18.2 terminator, f) pNOS:BlpR. The GoldenGate assembly technique was done as described in (Lampropoulos et al., 2013). Resulting pBLAX::XVE>>XXM-ER was transformed to GV3101 strain of *Agrobacterium tumefaciens* which was used for “floral dip” transformation of WT *Col-0*. T_1_ transformats were selected by BASTA resistance and strong estradiol induction of mCherry expression.

The GCaMP8s (Zhang et al., 2023) from pGP-CMV-jGCaMP8s vector (AddGene #162371) was BsaI domestificated by PCR amplification (primers: GCaMP_toC-F & GCaMP_mut-R; GCaMP_mut-F & GCaMP_toC-R) followed by NEBuilder ligation with BsaI restricted pGGC000 backbone. For assembly to pBLAX000 destination vector following blocks were combined: a) pUBQ10, b) N-decoy, c) GCaMP8s, d) C-decoy, e) HSP18.2 terminator, f) pNOS:HPT. Arabidopsis transformation and Hygromycin selection was done as described for XXM-ER.

XXM-ER_toC-F: gaagtgaagcttggtctcaggctccatgcgaccccaaatactcctc

XXM-ER_toC-R: tagggcgagaattcggtctcactgaggtggcggccgcgggtaccgc

ERret_F: ggcctgaaacgctattttctgaaaaaaaaactgatttaactgctgagaccgaattctcg

ERret_R: aatcagtttttttttcagaaaatagcgtttcaggcccttgtacagctcgtccatgcc

GCaMP_toC-F: gaagtgaagcttggtctcaggctcaatgcatcatcaccatcatcacacg

GCaMP_toC-R: gagaattcggtctcactgattacttcgctgtcatcatttgtac

GCaMP_mut-F: gggacggtgatgagatctctcggacac

GCaMP_mut-R: gtgtccgagagatctcatcaccgtcc

### Optogenetics

Constitutive XXM2.0 line was grown under spectrum with red and far-red light only. The XVE>>XXM-ER seeds were plated on AM+ supplemented with 100 µM estradiol (Sigma-Aldrich, E8875) and 2 µM all-trans retinal (Cayman Chemical, CAY18449-100). For imaging, the seedlings were placed to bottom-glass chamber under agar block. Imaging for growth was done on Nikon Ti2E microscope equipped with dark enclosure. Condenser used for bright feel imaging was covered by red filter foil (Lee Filters, #026) and blue light treatment was done by external blue LEDs (200 µmol·s^-1^·m^-2^). For R-GECO, GCaMP and HPTS imaging the LSM880 with Plan-Apochromat 10x objective was used similarly. During R-GECO imaging the roots were illuminated by green 561 nm excitation laser in both the light and dark phase. During the HPTS imaging the 488nm excitation laser served also as light source activating the XXM. As the imaging of dark phase is missing in the HPTS experiment, the signal was normalised to first frame, expecting the pH change to be minimal in first sec. of light stimulation. The GCaMP was imaged only 1/min to reduce exposure to low intensity (0.5%) 488nm excitation laser, the blue light treatment was done by external LEDs.

---

**Fig. S1.**
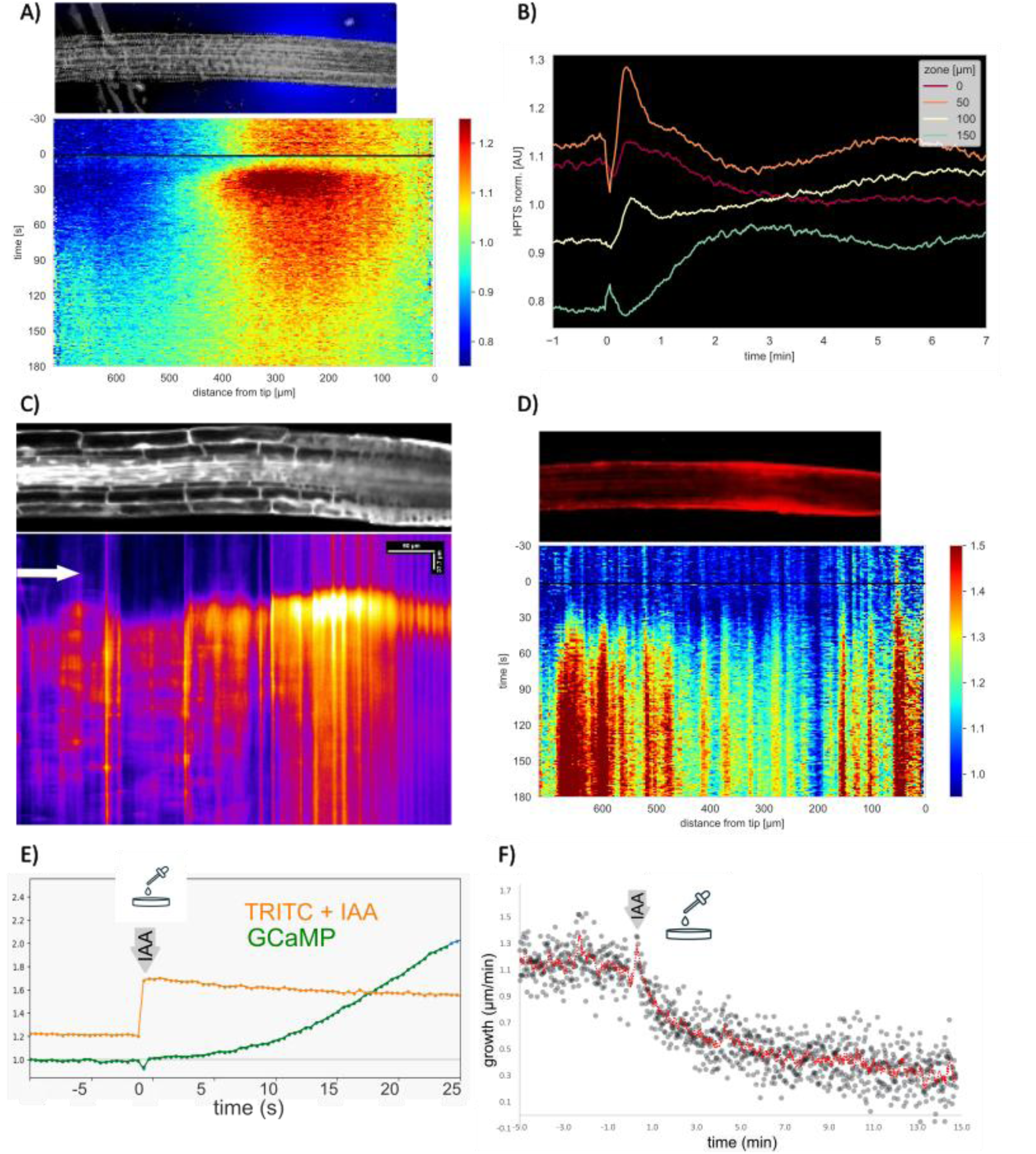
Spatiotemporal pattern of early IAA response in root. **(A)** Representative kymograph showing dynamics of HPTS halo (pH) in response to 100 nM IAA applied at t=0s. **(B)** Quantification of (A) The graphs represent mean signal in 50µm wide zones along the root tip. **(C)** Representative kymograph showing GCaMP response after 100nM IAA treatment. The signal increase is first visible in transition zone and then spreads. **(D)** Representative DiSBAC kymograph showing spatially uniform root tip depolarization upon 100 nM IAA **(E)** Sharp (<1 s) 100nM IAA treatment (arrow), confirmed by TRITC tracking dye mixed to the treatment cause increase of the cytosolic GCaMP (Ca^2+^) signal in <10s **(F)** and almost instant slowing of growth. The growth speed then decrees to ∼50%. (representative recording is shown)

**Fig. S2.**
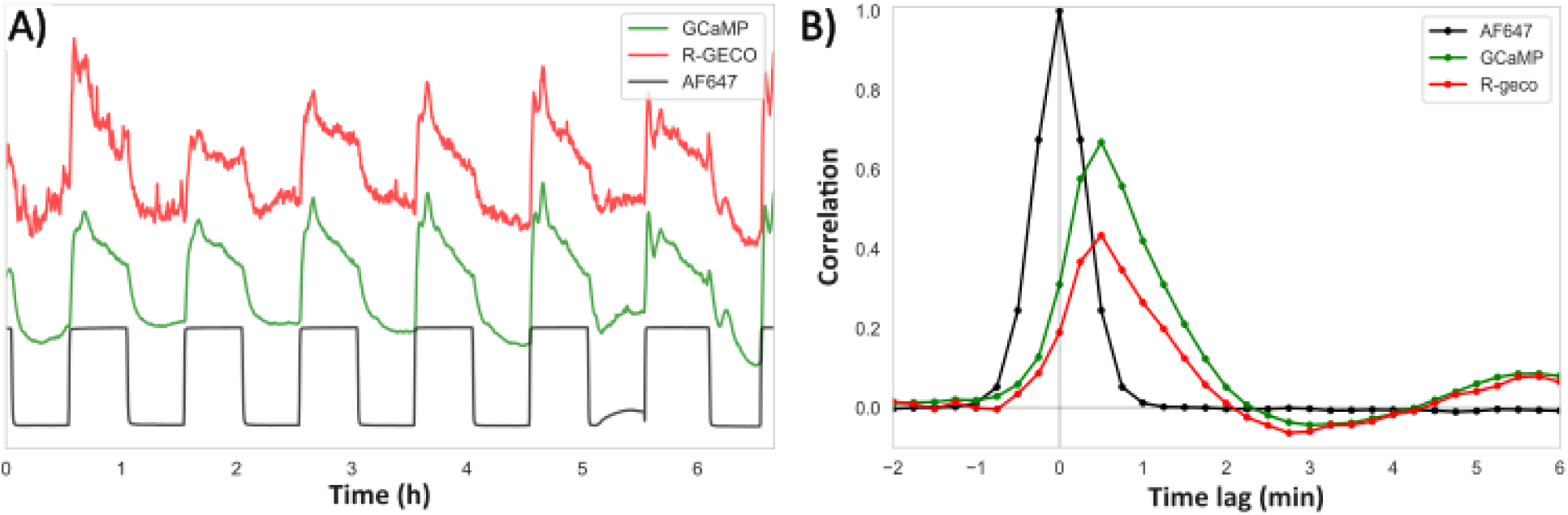
Calibrating the GCaMP and R-GECO response. **(A)** Signal from seedling co-expressing GCaMP and R-GECO during IAA oscillations (reported by AF^647^ tracking dye) in RootChip **(B)** Cross-correlation of signal of GCaMP and R-GECO to the AF^647^ reveals similar dynamic of both Ca^2+^ reporters.

**Fig. S3.**
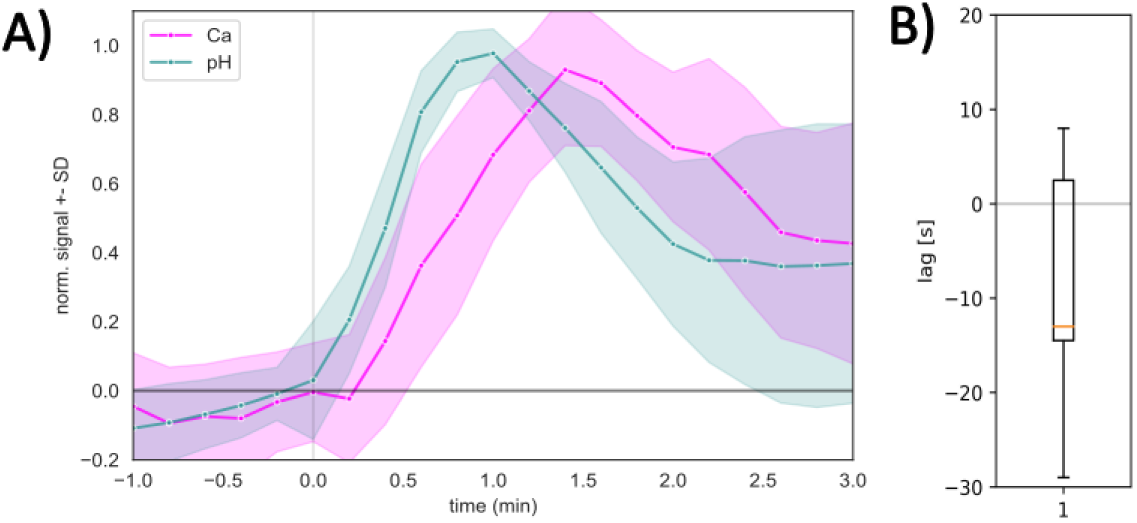
Cytosolic Ca^2+^ and apoplectic pH dynamic in transition zone. **(A)** HPTS (pH indicator) halo response precedes cytosolic R-GECO (Ca^2+^ indicator) in response to direct application of 100 nM IAA. Data represents the average across root transition zone. **(B)** The temporal lag determined by cross-correlation analysis of signals shown in (A).

**Fig. S4.**
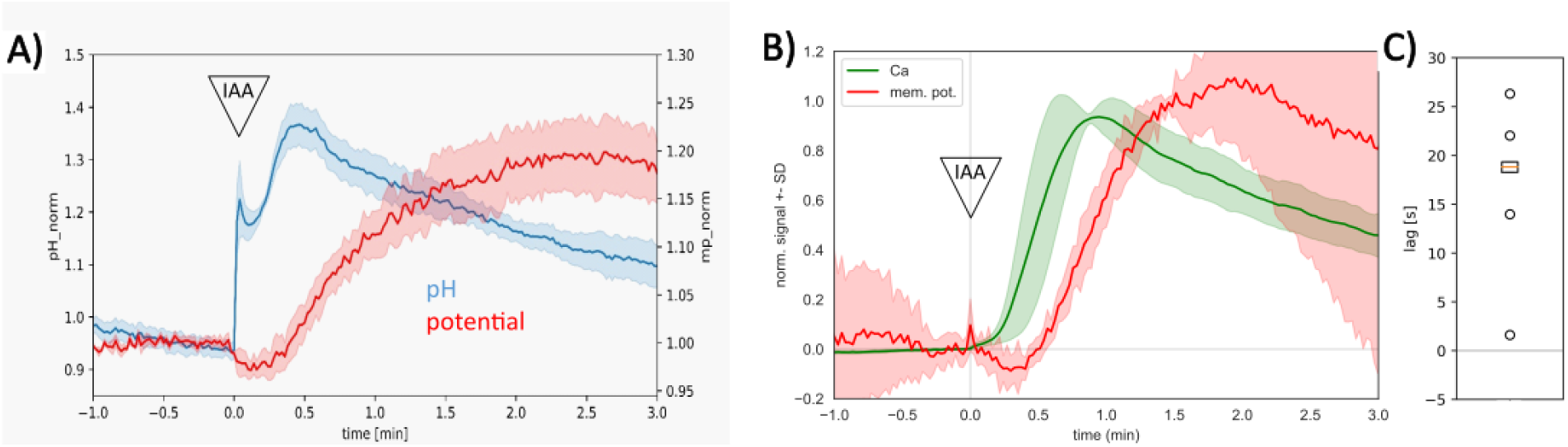
Apoplastic pH and cytosolic Ca^2+^ precede DiSBAC-reported depolarization. **(A)** Apoplastic HPTS (pH indicator) response precedes DiSBAC Signal. Data represents the average across whole root tip. The initial sharp increase in the HPTS signal is an artefact of the treatment application. **(B)** Cytosolic GCaMP (Ca^2+^ indicator) response precedes DiSBAC Signal. Data represents the average across whole root tip. **(C)** The temporal lag determined by cross-correlation analysis of signal pairs shown in (B).

**Fig. S5.**
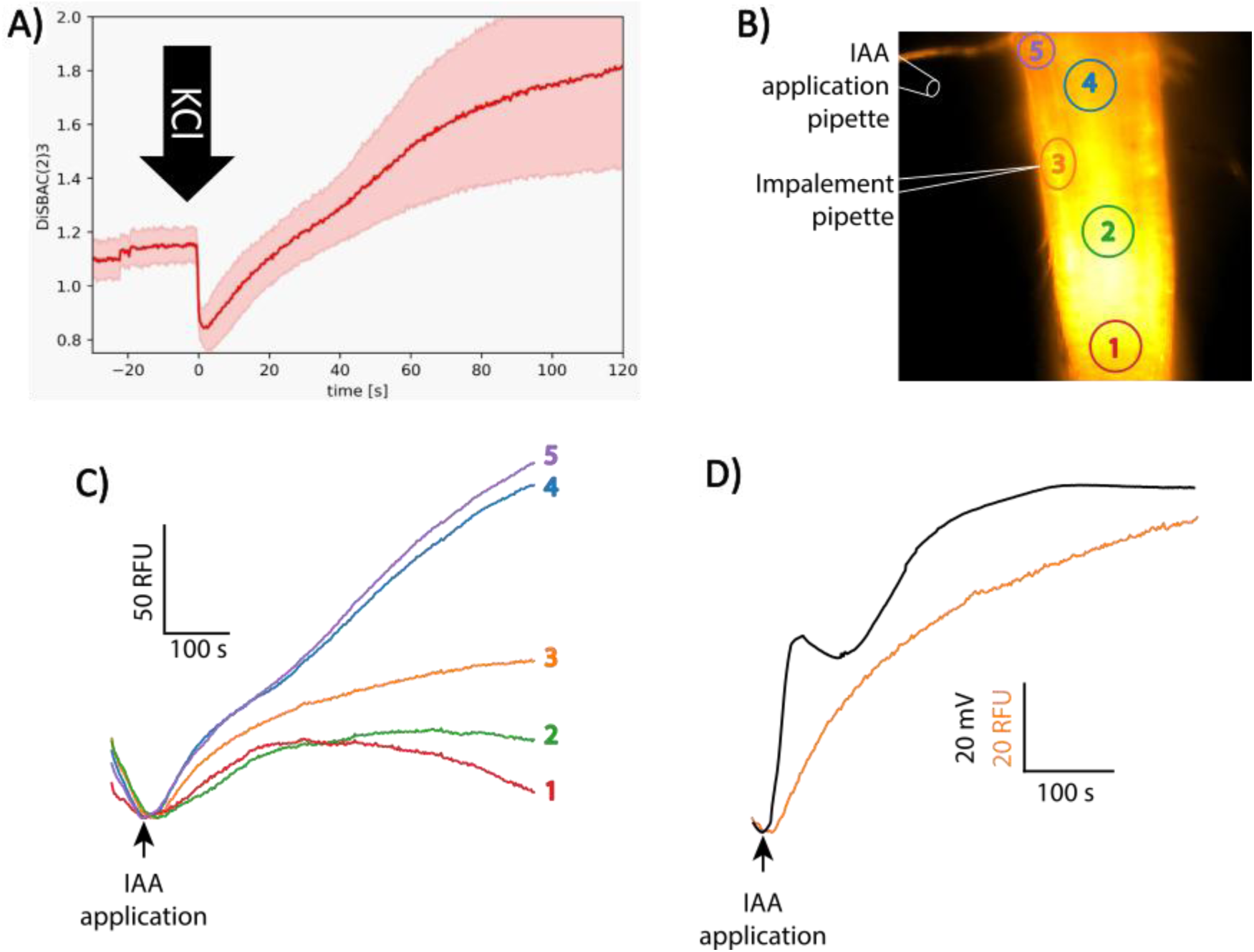
Calibrating the response time of DiSBAC. DiSBAC is typically considered a “slow” membrane potential dye, making it unsuitable for millisecond (ms) scale neurobiological recordings. To confirm whether DiSBAC is suitable for second (sec) scale timing in roots, **(A)** seedlings were treated with AM+ supplemented with additional 30 mM KCl, which is known to depolarize the plasma membrane. The initial brief drop in the DiSBAC signal was followed by a rapid rise, confirming the dye’s fast enough response for sec-resolution timing. Not a weak drop in the DiSBAC signal observed also after IAA application (sup. fig 2). **(B)** Setup for simultaneous recordings of membrane potential using the DiSBAC dye (orange) and the microelectrode. DiSBAC intensity was quantified in the numbered regions. **(C)** Following the application of 10 µM IAA the DiSBAC signal begins to rise almost immediately. **(D)** The initial rise of the DiSBAC signal overlaps with the microelectrode recording, however later, the DiSBAC signal rises more slowly than the electrode trace. Performed tests (A,C), confirm DiSBAC as a suitable method for timing membrane potential changes with second-scale resolution in roots.

**Fig. S6.**
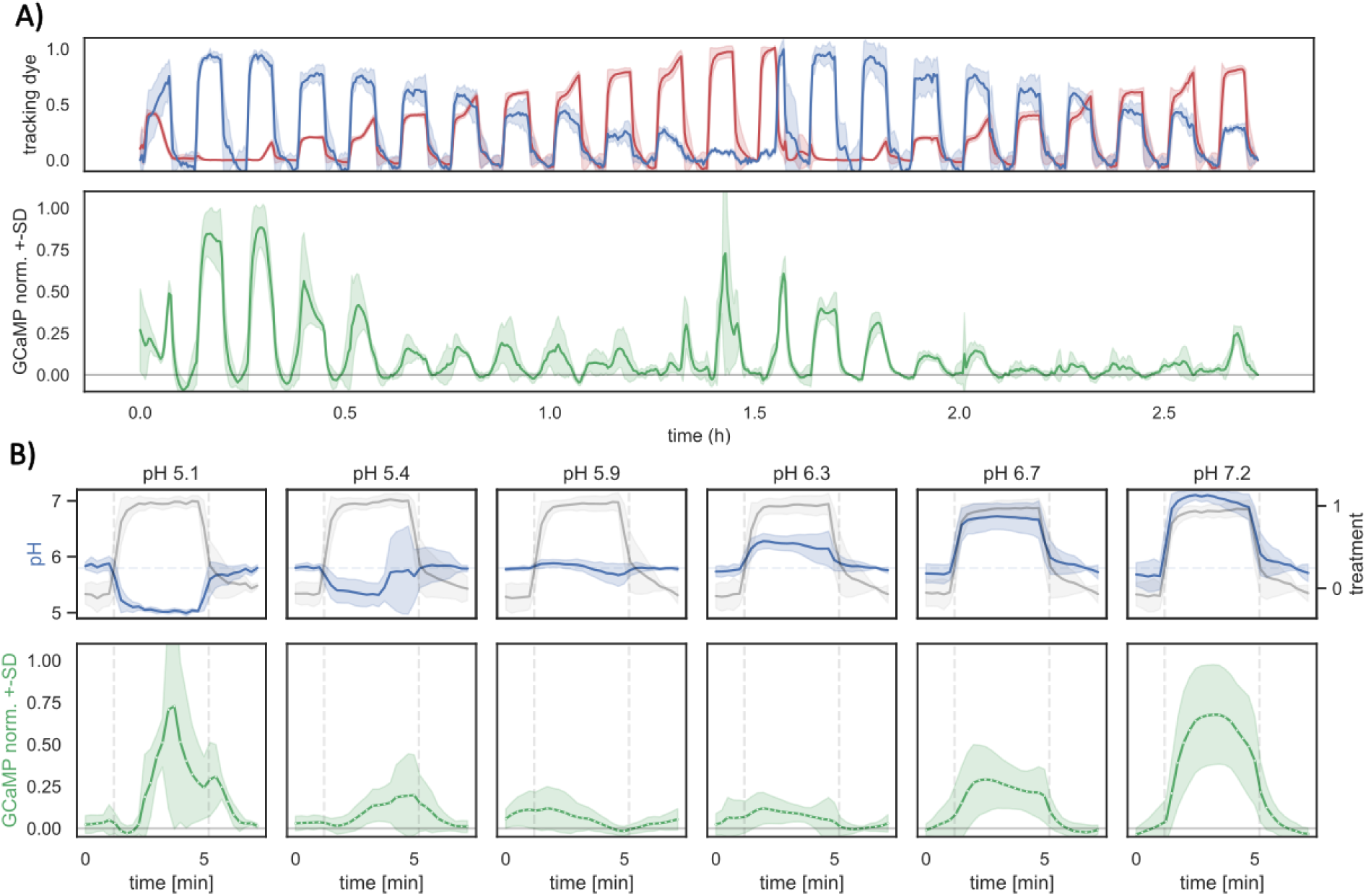
Dynamic of GCaMP response to the change of medium pH. **(A)** In the 3-inlet RootChip (Fig. 2C,E), repeated cycles of injection were performed using two types of media: a standard AM+ medium (pH 5.8, no tracking dye) and a mixed solution. The mixture was composed of 2 stocks in variant ratios: acidic AM+ (pH 5, labelled with TRITC) and alkaline AM+ (pH 7.3, labelled with AF647). **(B)** The treatment cycles were binned based on pH of treatment (blue, the bin average in the graph heading). The treatments were performed consistently across cycles, as shown by mock / mixture medium exchange (black). While medium alkalinisation (pH 5.8>7.2) is leading to constant elevation of GCaMP signal, the acidification (pH 5.8>5.1) resulted in a delayed GCaMP spike.

**Fig. S7.**
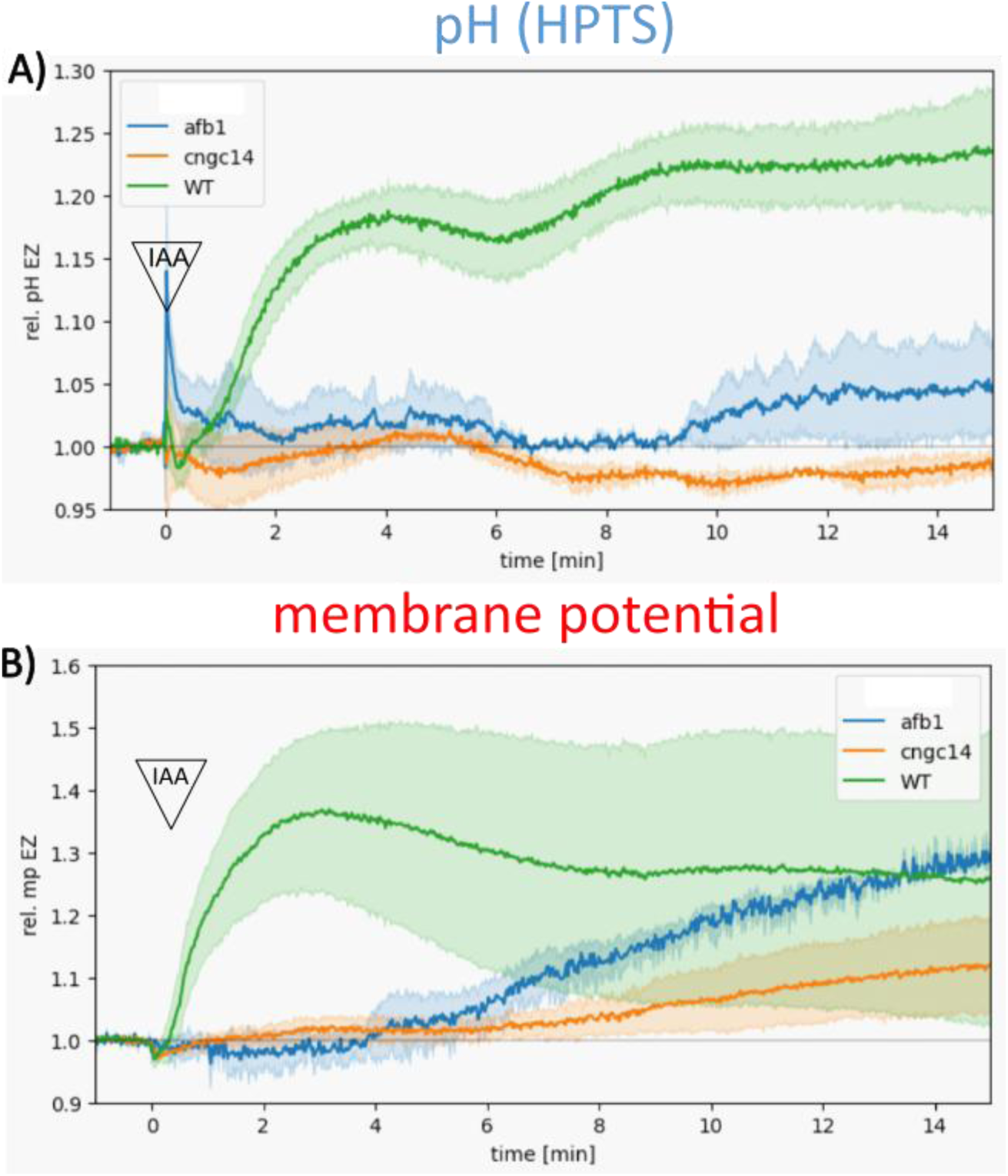
CNGC14 is required for fast IAA response. **(A)** The fast apoplastic pH response, monitored by HPTS, in the root elongation zone is insensitive to 100 nM IAA direct application in both the *cngc14* Ca^2+^ channel mutant and the *afb1* auxin receptor mutant. **(B)** The membrane potential response, monitored by the DiSBAC dye, is also insensitive to 100nM IAA in both the *cngc14* and *afb1* mutants

**Movie S1. Cytosolic Ca^2+^ and extraelular pH early IAA response**

Full representative movie from which selected stills are depicted as (Fig. 1B). Cytosolic R-GECO signal (Ca^2+^, magenta) and HPTS halo (pH, Cyan) shows distinct dynamic after direct application of 100 nM IAA treatment at t=0 min. (representative from 8 repetitions)

**Movie S2. Apoplast alkalinasation and PM depolarization has distinct dynamic**

Burst of HPTS halo (alkaline pH, blue) precedes gradual increase in DiSBAC signal (depolarisation of membrane potential, red) after direct application of 100 nM IAA treatment at t=0 min. (representative experiment)

**Movie S3. Lack of apoplastic Ca^2+^ cause root hair bursting**

Replacement of AM+ (1.5 mM Ca^2+^, tracked by TRITC dye, orange “grain”) to Ca^2+^-free medium (0.1 mM BAPTA, no “grain”) leads to acceleration of growth and bursting of root hairs. By re-introducing the Ca^2+^ medium, the growth slows down and new growing root hairs emerge.

